# Investigation of H9N2 avian influenza immune escape mutant that lacks haemagglutination activity

**DOI:** 10.1101/2023.10.03.558847

**Authors:** Thusitha K. Karunarathna, Jean-Remy Sadeyen, Sushant Bhat, Pengxiang Chang, Jiayun Yang, Mehnaz Qureshi, Joshua E. Sealy, Rebecca Daines, Munir Iqbal

## Abstract

H9N2 avian influenza viruses pose a global threat to animal and human health. While vaccination is essential for mitigating disease impact, these viruses evolve to evade vaccine immunity through changes in the haemagglutinin (HA) glycoprotein. In this study, we identified immune escape mutation in an H9N2 virus resulting from pressure exerted by homologous chicken antisera. The immune-escape variant acquired an amino acid substitution, replacing glycine (G) with glutamic acid (E) at position 149 in the HA protein. The G149E mutant virus lost the ability to agglutinate chicken erythrocytes, while still maintaining replication comparable to the wild-type virus in chicken embryos and cells. This led to the hypothesis that the G149E substitution, leading to a shift from a neutral to a negative charge polarity at HA position 149, might be crucial for the optimal interaction between the virus and receptors on erythrocytes. Investigation indicated that agglutination could be restored by substituting E to positively charged amino acids histidine (H), arginine (R) or lysine (K). These findings suggest that the H9N2 virus may be likely acquire the G149E mutation under immune pressure in nature. This mutation poses challenges to vaccination and surveillance efforts as it partially evades immune protection and is not easily detectable by conventional haemagglutination assays. This underscores the intricate interplay between antigenic variation and viral traits, emphasising the critical need for ongoing surveillance and research to effectively mitigate the risks associated with avian influenza H9N2 viruses.

**IMPORTANCE:** Understanding how avian influenza viruses evolve to persist in nature is crucial for enhancing disease mitigation tools such as vaccines, diagnostics, and risk assessment. In this study, we identified an H9N2 virus antibody escape mutant with G149E mutation in the haemagglutinin that had lost the ability to agglutinate chicken erythrocytes, while retaining infectivity and replication fitness. The lack of haemagglutination activity potentially negatively impacts routine surveillance and commonly used diagnostics such as haemagglutination assay or haemagglutination inhibition assay. Therefore, it is urgent to develop and adopt alternative methods for viral detection. Difficult to detect variants potentially that are not compatible with common surveillance techniques could circulate remain silent while reassort with other influenza viruses, which posing unpredictable risks to animal and human health. This research helps us better understand avian influenza, leading to improved disease control, diagnostics, and risk assessment to protect both animals and humans.

## Introduction

Avian Influenza viruses (AIVs) belong to *Orthomyxoviridae*, which are segmented, negative-sense RNA viruses. AIVs pose a major threat to global food security (1) and can cause zoonotic infection in humans, and are considered viruses with pandemic potential (2, 3). H9N2 is one of the widely prevalent AIV subtypes in poultry and remains enzootic in many countries worldwide (4–7). H9N2 AIVs are classified as low pathogenicity in chickens compared to the high pathogenicity subtypes (polybasic site containing H5 and H7) (5).

In recent years, H9N2 viruses have acquired genetic changes that increase their systemic tropism, resulting in severe morbidity and increased mortality rate (up to 60%) (8). Additionally, H9N2-infected chickens experience greater morbidity and mortality due to secondary bacterial infections or co-infections with other avian viruses (9, 10). The transmission of H9N2 AIVs occurs through respiratory droplets and fomites, including feed and water contamination (11, 12). Recent studies indicate that the contemporary H9N2 AIVs prevalent in poultry have acquired genetic changes in the haemagglutinin (HA) gene, which have widened viral receptor binding affinity to both avian and human cell surface receptors and enhanced zoonotic risks (13). This has been supported by the highest reported human infections (>60) in the past three years in regions where H9N2 is endemic in poultry (2, 14-16). H9N2 viruses are also very efficient in donating internal gene segments to other AIVs (17, 18). This genetic reassortment mechanism has led to the emergence of several highly zoonotic AIVs, including H5N1(4), H7N9 (19), H10N8 (20), H5N6 (21), and H3N8 (22).

The hyper-prevalence of H9N2 results in huge economic losses in many countries, which has led the governments to adopt vaccination programs administering adjuvanted, inactivated virus vaccines. The neutralizing antibodies generated against the virus’s hemagglutinin antigen following vaccination or infection directly inhibit the virus’s attachment to the target receptor molecules present on the host cell surface (23). Thus, to overcome antibody-based neutralisation, the virus evolves using an immune escape mechanism (24–26). This has led to the emergence of immune escape variants that carry amino acid changes in their HA genes. The genotypic changes often lead to viral phenotypic change, such as infectivity, replication efficiency, viral fusion, thermal stability, receptor binding, and/or hemagglutination activity (27, 28). Therefore, it is critical to closely monitor virus evolution to identify emerging variants in the field. Haemagglutination assay is routinely used as a detection method for virus isolation from field samples and haemagglutination inhibition (HI) assay are widely used for antigenic characterization (29, 30). In recent years antigenic changes in human H1N1 and H3N2 viruses failed to agglutinate erythrocytes, which caused the viruses are undetectable by haemagglutination assay and HI assay (31–34). It is possible that such failure of detection occurs in other influenza subtypes. This will pose challenges to virus isolation and antigenic characterization. (31, 35).

In this study, we conducted an immune escape experiment to identify mutants of the H9N2 strain (A/chicken/Pakistan/UDL-01/2008, hereafter referred to as wt-UDL-01/08) that can overcome the neutralizing activity of antibodies in the sera from vaccinated chickens. We successfully identified an immune escape variant carrying the G149E substitution in the HA gene. Interestingly, this G149E mutant (hereafter referred as mutant G149E-UDL-01/08 virus) was unable to agglutinate chicken erythrocytes. Through this analysis, we sought to define the underlying mechanisms that potentially contribute to the loss of agglutination activity observed in this mutant. Our findings from this investigation provide valuable insights into the risks of epizootic and zoonotic threats posed by emerging H9N2 variants that may lack agglutination activity and persist undetected in the global poultry population, highlighting the importance of continued surveillance and monitoring efforts to safeguard public health and the poultry industry.

## Results

### The G149E mutation on the H9HA of the wt-UDL-01/08 virus results in a loss of agglutination of chicken erythrocytes

We conducted immune escape experiments (manuscript is in preparation) and identified G149E substitution in HA (based on mature H9 peptide numbering) that lacked haemagglutination activity (Supplementary Figure 1). While there have been reports describing human influenza viruses that could not agglutinate chicken red blood cells (cRBC) (32, 34), we were unable to find any previous reports on loss of agglutination activity in H9N2 strains. The propagation of this mutant G149E-UDL-01/08 virus in embryonated chicken eggs showed 10-fold higher end-point titres compared to the progenitor wt-UDL-01/08 virus (Table 1). Altogether, the results suggests that despite a higher PFU/ml virus input of mutant G149E-UDL-01/08 virus, we were still unable to detect the HA activity, indicating that the G149E mutation in H9HA in UDL-01/08 virus results in the loss its haemagglutination activity.

**Table 1.** Haemagglutination and Virus titres of wt-UDL-01/08 and mutant G149E-UDL-01/08 viruses.

| Name of virus | Virus titre (PFU/ml) | HA titre<sup>a</sup> (HAU/50µl) |
| --- | --- | --- |
| wt-UDL1/08 | 5.00E+06 | 128 HAU/50µl |
| G149E-UDL-01/08 mutant | 4.75E+07 | < 2 HAU/50µl |

The haemagglutination (HA) titres are indicated as haemagglutination units (HAU) and virus titres are indicated as plaque forming units (PFU/ml). The virus titres were determined by plaque assay in MDCK cells. The HAU were determined using chicken RBCs and viruses were grown in chicken embryonated eggs. <sup>a</sup>Titres are expressed as the reciprocal of the highest virus dilution producing haemagglutination. G (glycine) and E (glutamic acid).

### The G149E mutant virus showed reduced antigenic cross-reactivity against the wt-UDL-01/08 virus

To evaluate whether the G149E mutation in the H9HA altered the antigenic properties of the wt-UDL-01/08 virus, we performed a microneutralization (MNT) assay using polyclonal antisera (post-vaccination) raised against the wt-UDL-01/08 virus in a group of five chickens. The mutant G149E-UDL-01/08 virus showed up to a two-fold reduction in MNT titers compared to the wt-UDL-01/08 virus (Figure 1). The observed reduction in MNT titers in serum samples was statistically significant (P< 0.0039). Suggesting that the emergence of variants with G149E mutations may also compromise the effectiveness of vaccine.

**Figure 1.**
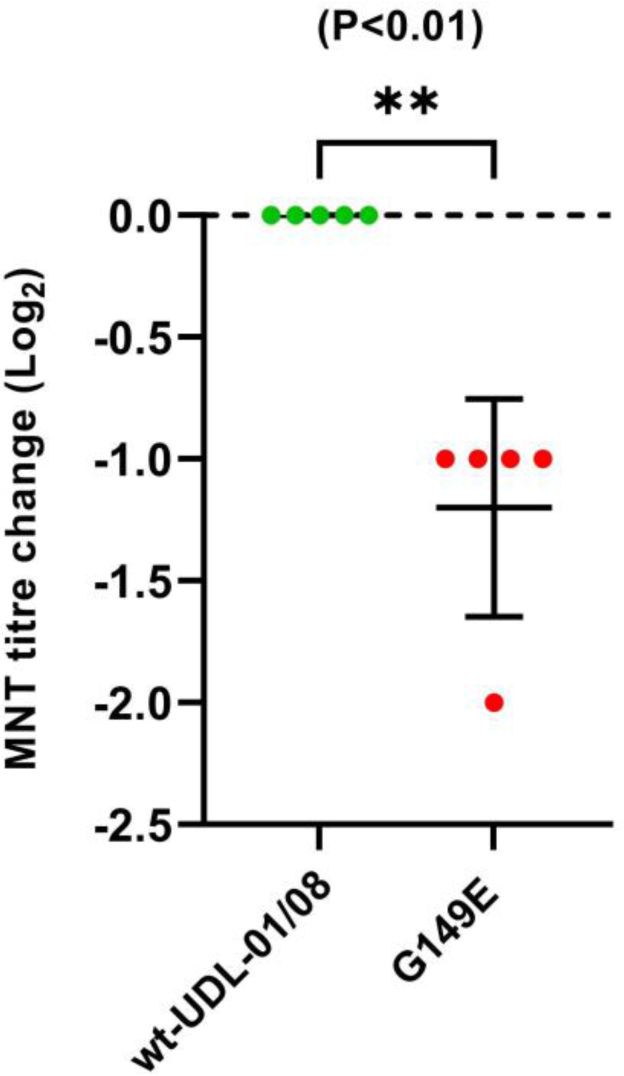
Microneutralization (MNT) titre change of G149E-UDL-01/08 mutant virus against post-vaccination chicken sera raised against wt-UDL-01/08 viruses. MNT assays were carried out with a panel of 5 post-vaccinated chicken antisera raised against the wt-UDL-01/08 virus and each dot represents the MNT titre from an individual chicken serum sample. MNT titres of each serum samples against wt-UDL-01/08 virus were normalized to 0 and used to compare the MNT titre change of mutant viruses. Paired t-test was used to compare MNT titre of wt-UDL-01/08 virus to G149E-UDL-01/08 mutant virus (** p<0.0039). SD of the mean shown.

### Loss of chicken erythrocytes agglutination is a strain specific phenomenon

To investigate the loss of agglutination to cRBCs with the G149E mutation is specific to UDL-01/08 or universal for other H9N2 lineages, we generated two additional mutant viruses harbouring G149E mutation in their HA genes. One mutant G149E-A/Ck/Vietnam/315/2015 virus (G149E-VN/315/15), representing the BJ/94 lineage, predominant in China and Vietnam, and the other mutant G149E-A/Env/Bangladesh/26218/2015 (G149E-BD/26218/15) representing the same G1 lineage as UDL1/08. Both H9N2 strains (G149E-VN/315/15 and G149E-BD/26218/15) with G149E mutation showed comparable haemagglutination activity in comparison to corresponding wild-type (wt) viruses (Table 2). Therefore, the loss of haemagglutination activity by G149E mutation in the HA is not consistently observed across all strains of H9N2 viruses, including within the G1 lineage.

**Table 2.** Haemagglutination and virus titres of wild-type viruses (wt-VN/315/15 and wt-BD/26218/15) and their mutant (G149E-VN/315/15 and G149E-BD/26218/15) viruses.

| Name of virus | Virus titre (PFU/ml) | HA titre<sup>a</sup> (HAU/50µl) |
| --- | --- | --- |
| wt-VN/315/15 | 2.00E+08 | 256 HAU/50µl |
| G149E-VN/315/15 | 3.50E+07 | 128 HAU/50µl |
| wt-BD/26218/15 | 6.00E+06 | 64 HAU/50µl |
| G149E- BD/26218/15 | 7.25E+06 | 64 HAU/50µl |

### The specific composition of the Receptor Binding Site (RBS) in HA genes is critical for haemagglutination activity of H9N2 viruses

The receptor binding site (RBS) of HA is comprised of three secondary structure elements: the 130-loop, the 190-helix, and the 220-loop. The RBS is mainly involved in receptor binding activity, binding to sialic acid receptor on cells of target tissues, including cRBC (36). As we observed, the G149E mutation in the wt-UDL-01/08 virus abolished agglutination activity, but the same mutation had no effect on the agglutination activity of the other two H9N2 viruses, VN/315/15 and BD/26218/15 (Table 2). Therefore, we analysed molecular differences in the RBS between wt-UDL-01/08 and wt-BD/26218/15 viruses, which are both G1 lineage. Six amino acids differences were identified, three of which are located in the 190-helix and the other three amino acids are within the 220-loop at the RBS (Figure 2).

**Figure 2.**
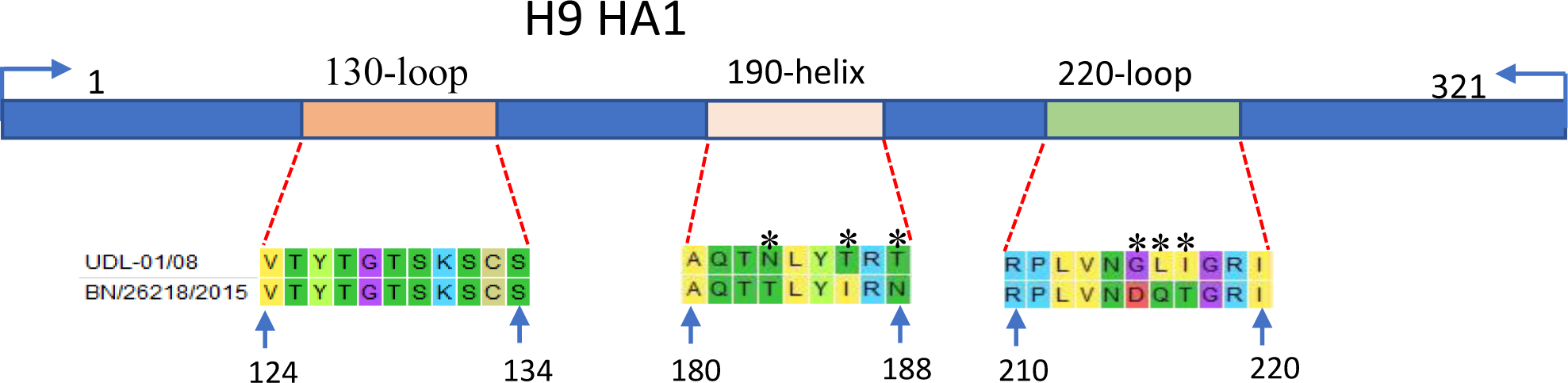
Alignment of RBS of HA1 region in HA gene of wt-UDL-01/08 and wt-BD/26218/15 viruses. Difference in amino acid sequences have been indicated in asterisk at position 183, 186, 188 (in 190-helix) and 215, 216 and 217 (220-loope).

To further analyse the key residues responsible for the agglutination activity of the wt-UDL-01/08 virus, we generated two variants of the mutant G149E-UDL-01/08 virus. These variants contained either the 190-helix or 220-loop of the RBS from the BD/26218/15-like virus. The haemagglutination assays of these viruses showed that when we substituted three amino acid residues (G215D, L216Q, I217T) from the 220-loop of the BD/26218/15 virus into the G149E-UDL-1/08 mutant virus, it regained its haemagglutination activity. However, the G149E-UDL-1/08 mutant containing the 190-helix similar to the BD/26218/15 virus did not regain the ability of haemagglutination (Table 3 and Supplementary Figure 2). These results suggest that, when combined with the G149E mutation, the composition of the 220 loop in the wt-UDL-01/08 virus contributes to the loss of its haemagglutination activity.

**Table 3.** Haemagglutination and virus titres of mutant viruses of UDL-01/08 virus carrying G149E with converted their 190-helix and 220-loop of RBS separately into BD/26218/15 virus’ RBS.

| Name of virus | Virus titre<br>(PFU/ml) | HA titre<br>(HAU/50µl) |
| --- | --- | --- |
| G149E-UDL-01/08 HA-N183T+T186I+T188N | 1.50E+05 | 0 |
| G149E-UDL-01/08 HA-G215D+L216Q+I217T | 1.50E+05 | 4 |
| wt-UDL1/08 | 5.00E+06 | 128 |

To further validate the significance of each of these residues in contributing to the agglutination activity of H9N2 viruses, we generated additional mutants of G149E-UDL-01/08 virus containing G215D, L216Q, or I217T substitutions. Through haemagglutination assays, we confirmed that amino acids D, Q or T at positions 215, 216, and 217 in the HA of H9N2 viruses play a crucial role in maintaining the haemagglutination activity (Supplementary Figure 3 and Table 4).

**Table 4.**
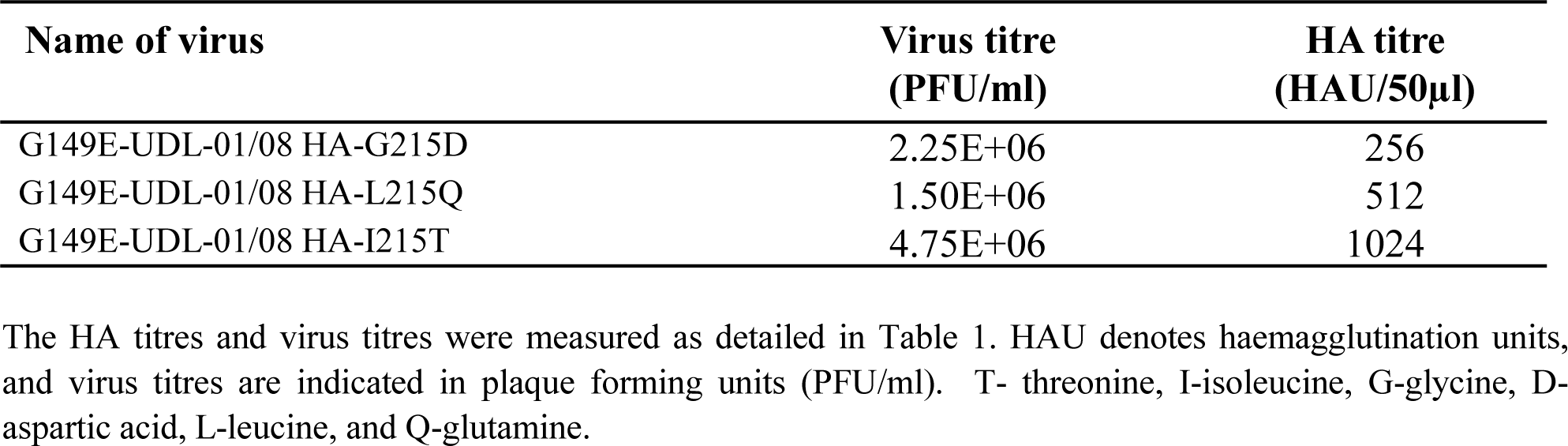
Haemagglutination titres with chicken erythrocytes and virus titres (PFU/ml) of mutant viruses of UDL-01/08 virus carrying G149E with converted position at 215, 216, 217 in 220-loop of RBS separately into BD/26218/15 virus’ RBS.

### The polarity of the amino acid at position 149 in the H9 haemagglutinin regulates the virus haemagglutination activity

Sialic acid, and in particular sulphated sialylated glycans that are the preferential receptor for H9N2 viruses such as UDL-01/08 (37) and negatively charged. We have previously found receptor avidity of H9N2 viruses to sulphated receptors correlate crudely with net charge of the HA1 (38). We hypothesised that changes in the amino acid composition at position 149 might be linked to alterations in the HA charge, crucial for maintaining an optimal interaction between the virus and the cellular receptor on RBCs. This balance is likely critical for viral haemagglutination activity. To determine whether a shift in the polarity of the amino acid at position 149 is associated with a loss of haemagglutination activity, we generated mutant UDL-01/08 viruses carrying either positively charged (G149K, G149R, and G149H) or negatively charged (G149D) residues at HA position 149. The mutant viruses with amino acid "D" at residue 149 in the HA showed no haemagglutination of chicken erythrocytes, similar to G149E. Conversely, the mutants carrying positively charged amino acids "K, R, or H" at HA position 149 retained their haemagglutination activity. Interestingly, the polarity shift did not impact the replication profiles of these mutant viruses. The mutant virus containing the negatively charged amino acid (G149D) that lost its haemagglutination activity not only retained its replication fitness but also exhibited a more than two-fold increase in virus end point titres when propagated in embryonated chicken eggs compared to the mutants with positively charged amino acids (G149K, G149R, and G149H) (Table 5). These findings suggest that positive polarity at the HA residue 149 is crucial for the agglutination of chicken erythrocytes by the H9N2 wt-UDL-01/08 virus.

**Table 5.** Haemagglutination titres and virus titres (PFU/ml) of mutant viruses of UDL-01/08 processing HA mutations at 149 with negatively charged (E-glutamic acid, D-aspartic acid) and positively charged (K-Lysine, H-Histidine, R-Arginine) amino acids with chicken erythrocytes.

| Name of virus | Virus titre (PFU/ml) | HA titre<sup>a</sup> |
| --- | --- | --- |
| UDL-01/08-G149E | 4.75E+07 | <2 HAU/50µl |
| UDL-01/08-G149D | 2.25E+07 | <2 HAU/50µl |
| UDL-01/08-G149K | 8.75E+06 | 128 HAU/50µl |
| UDL-01/08-G149H | 7.50E+06 | 256 HAU/50µl |
| UDL-01/08-G149R | 3.25E+06 | 128 HAU/50µl |

The HA structure prediction analysis also showed that substitution of amino acid from “G” to “E” at HA 149 position increase the negative charge distribution in and around the HA receptor binding site due to the introduction of the carboxyl group from glutamic acid (Supplementary Figure 4B). Conversely, the HA with residues “K” at position 149 displays a reduction in negative charge distribution in the receptor binding site because the carboxyl group from glutamic acid is replaced by an amino group from lysine (Supplementary Figure 4C). This overall change in charge distribution in the vicinity of the receptor binding site may explain the distinct phenotypic changes between the G149E and G149K viruses as depicted in our proposed model (Figure 6).

**Figure 6.**
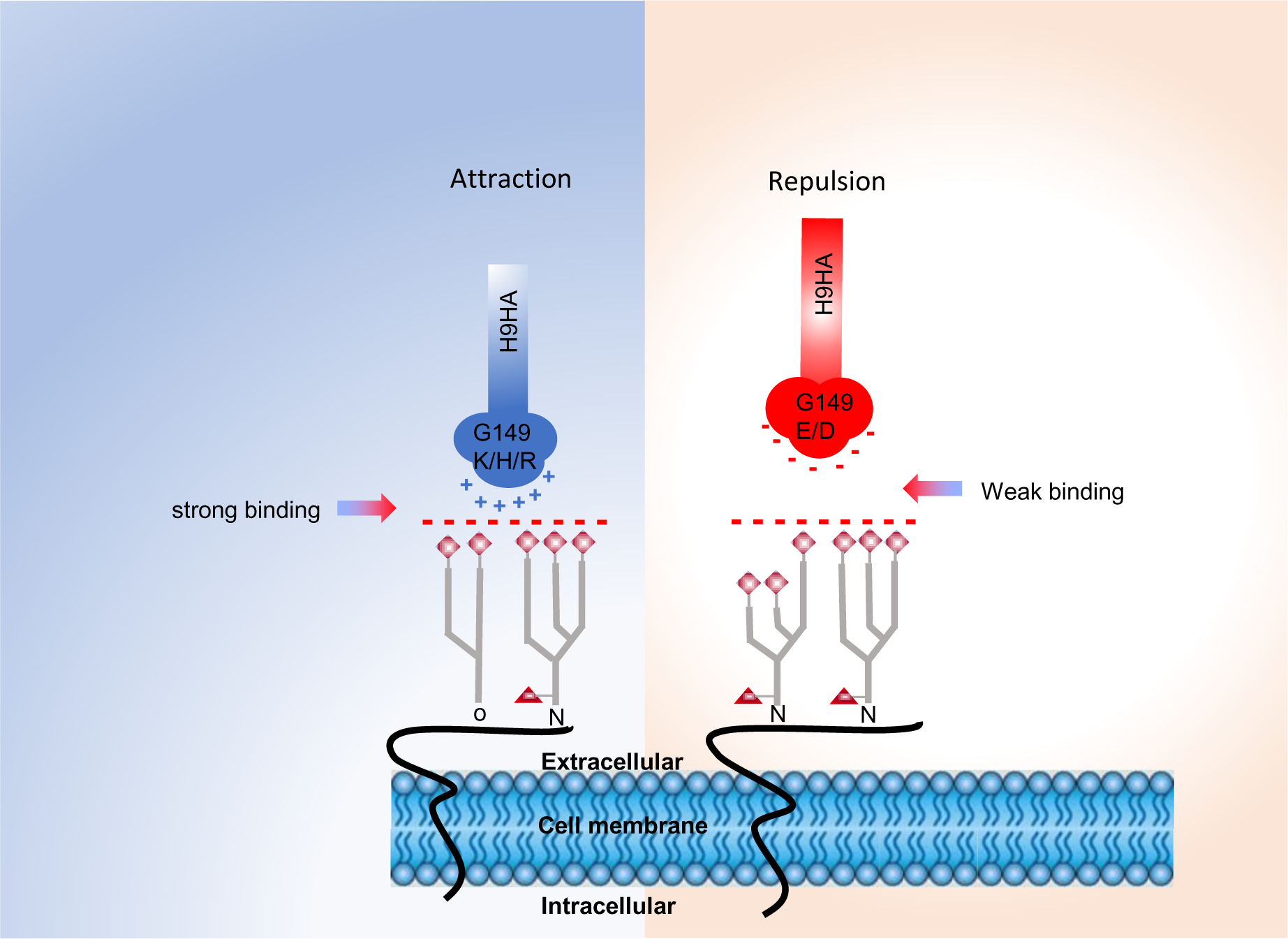
Graphical presentation of the proposed model for loss of the ability to agglutinate chicken erythrocytes with negatively charged H9HA. The mutant viruses carrying negatively charged amino acid substitutions at position 149 (G149E and G149D) exhibit an increased in the negative charge distribution (indicated in red colour on the HA protein), mainly in and around the receptor binding site due to the introduction of the carboxyl group from glutamic acid (E) and aspartic acid (D). Negatively charged O-linked and N-linked (indicated as O and N, respectively) sialic acid receptors are anchored to the cell membrane extracellularly. The negative charged sialic acids are indicated in red-coloured squares and dashes. The shift in charge distribution in H9HA towards a negative charge promotes weaker binding between chicken erythrocytes and virus particles, results in no agglutination. The mutant virus carrying positive amino acids (K-lysine, H-histidine, and R-arginine) increases the positive charge of H9HA (indicated in blue-coloured HA protein) and exhibits strong binding of virus particles to chicken erythrocytes, promoting haemagglutination. “+” indicates a positive charge, and “-” indicates a negative charge.

### The G149E mutant virus retained stability following multiple passage and with multiple H9N2 genetic backgrounds

Prior studies have noted that evolutionary changes in the HA that enable immune evasion are often followed by compensatory changes in HA or NA which restore the fitness cost that may have been incurred by the initial substitution (39, 40). To verify the stability of the G149E substitution, the mutant G149E-UDL-01/08 virus and the wt-UDL-01/08 virus, which served as a control, were passaged five times in embryonated chicken eggs. After each passage, the HA and NA coding regions of the progeny viruses were sequenced. The results revealed no additional synonymous or non-synonymous mutations, thereby validating that the G149E substitution maintained its stability within the mutant G149E-UDL-01/08 virus throughout all five passages.

To further assess the compatibility of the G149E substitution in the HA of various H9N2 viruses, two distantly related H9N2 viruses (Viet/315/2015 HA and BD/26218/2015) containing the G149E substitution were generated using reverse genetics (RG) approach. Both viruses were successfully rescued on the first attempt. These mutant viruses demonstrated that the replication competence in embryonated eggs were similar to their wt counterparts. Additionally, sequence analysis confirmed that no substitutions had occurred in the HA and NA coding regions of these rescued viruses (data not shown). This suggests that the G149E substitution can maintain its fitness across different strains of H9N2 viruses.

### The G149E mutant virus showed reduced receptor binding avidity for both the avian-like **_α_**2,3-linked sialic acid analogue and human-like **_α_**2,6-linked sialic acid analogue

Changes in amino acid residues within or near the RBS can modulate the H9N2 virus’s receptor binding preferences, thereby influencing virus attachment and entry into host cells during infection (36). To assess the influence of the G149E mutation on the receptor binding profiles, a biolayer interferometry approach was employed for the G149E-UDL-01/08 mutant virus (33). Compared to the wt-UDL-01/08 virus, the mutant G149E-UDL-01/08 virus exhibited a 10-fold decrease in binding to the avian-like receptor Neu5Ac α-2,3Gal β1-4(6-HSO_3_) GlcNAc (3SLN(6su)). Furthermore, the mutant G149E-UDL-01/08 virus lost any detectable binding to human-like receptor analogue α2,6-sialyllactosamine (6SLN) (Figure 3). This suggests that the G149E mutation has reduced the binding affinity for both avian-like 3SLN(6su) and human-like 6SLN receptors compared to the wt-UDL-01/08.

**Figure 3.**
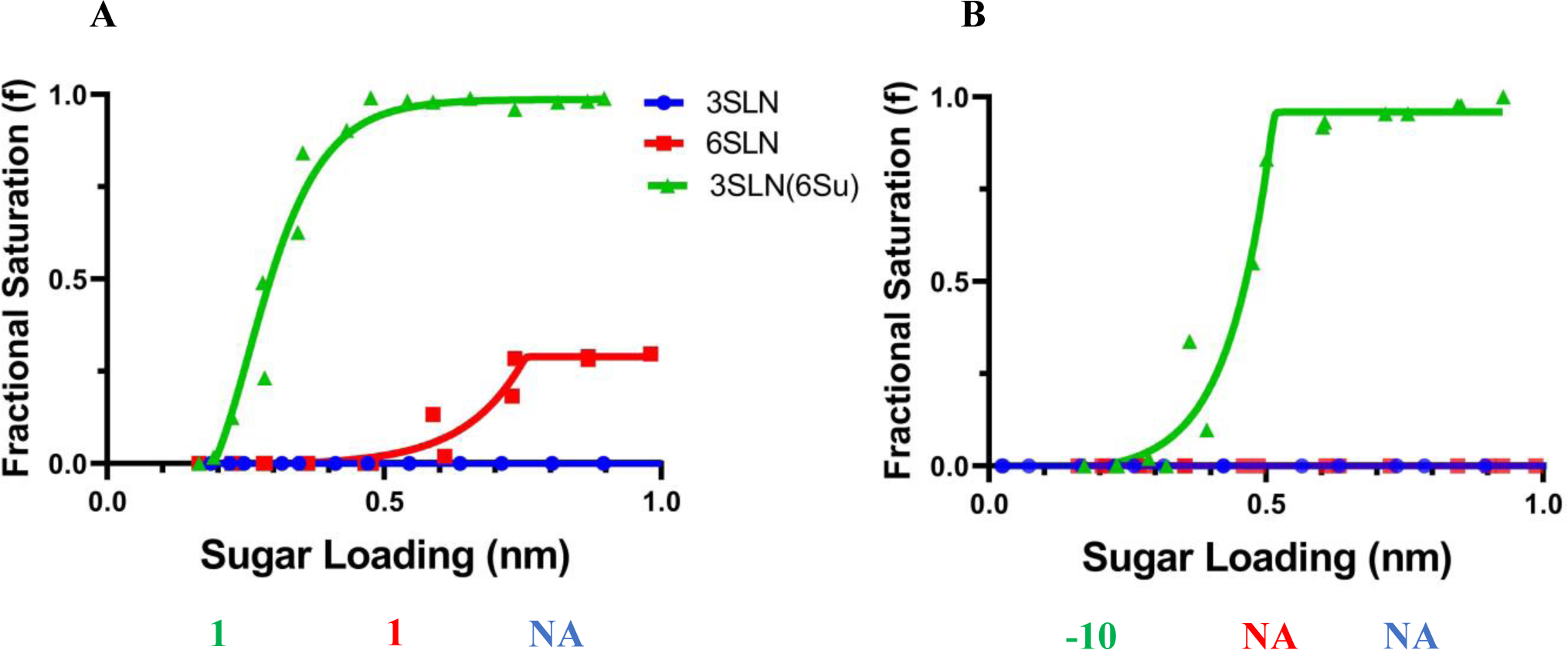
Characterization of the receptor binding properties of wt-UDL-01/08 and G149E-UDL-01/08 mutant viruses. The binding of purified wt-UDL-01/08 (A) and G149E-UDL-01/08 mutant (B) viruses to three different influenza receptor analogues was assayed by biolayer interferometry, blue lines shows α2,3 linked sialyllactosamine (SLN) (3SLN), red lines shows α2,6 linked SLN (6SLN) and green line shows α2,3 linked SLN with an additional sulphate residue on the 6’ position of the N-acetylglucosamine (3SLN(6Su)). All data is modelled with sigmoidal dose response curves and the amalgamation of two repeats. The numbers below each figure show the fold change of receptor binding of indicated viruses to α2,3 SLN (shown in blue), α2,6 SLN (shown in red) and α2,3SLN(6Su) (shown in green) receptor analogues compared to wt-UDL-01/08. -, reduction; +, increase; NA, not applicable.

### The G149E mutant virus retained replicative fitness comparable to the wt-type virus *in vitro* **and *in ovo*.**

To assess the impact of the G149E mutation on virus replication efficiency, both the mutant G149E-UDL-01/08 virus and the wt-UDL-01/08 viruses were assessed for their replication in MDCK cells (serving as a mammalian host model), primary chicken kidney cells (CKCs) and ten-day-old embryonated chickens’ eggs (serving as avian host models). Two additional mutants, G149D and G149K, were included as controls. These contained negatively and positively charged amino acids at the HA 149 position, respectively. The replication efficiency of all three mutant viruses in MDCK cells was similar to the wt-UDL-01/08 virus up to 12 hours. However, the mutant virus with the G149D mutation replicated at a significantly higher rate, showing 1.8-fold increase in viral titres compared to the wt-UDL-01/08 virus at 24 hours post-infection (hpi). At 48 hpi post-infection, the viral titres of both mutant viruses with a negatively charged amino acid at HA position 149 (G149E and G149D) was also significantly higher (2 and 2.6-fold respectively) compared to the wt-UDL-01/08 virus. In contrast, the mutant virus with the positively charged amino acid (G149K) in the HA replicated at a notably reduced rate, and its viral titres were comparatively lower at the same time points. Beyond 48 hpi, a consistent differential replication trend persisted between the viruses carrying the negatively charged and the positively charged amino acids at HA position 149. However, the differences in viral titres at this time point were not statistically significant (Figure 4A).

**Figure 4.**
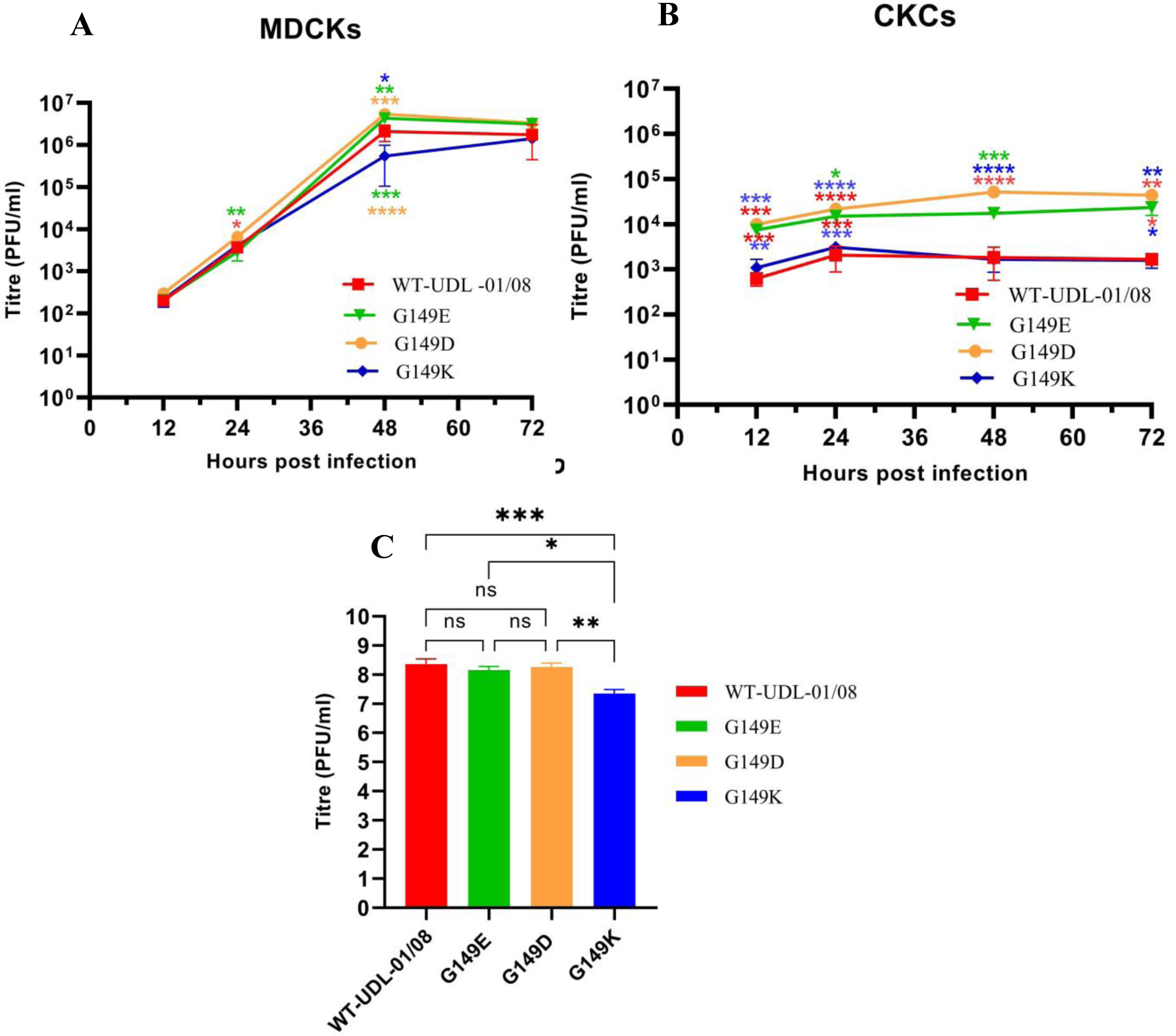
Virus replication kinetic of wt-UDL-01/08 and its mutant viruses carrying positively charged and negatively charged amino acid at passion 149 *in vitro* in MDCK (A), CK (B) cells and *in ovo* (C) MDCK and CKCs were infected in triplicate with a multiplicity of infection (MOI) of 0.001 and 0.01 respectively and supernatants were collected at 12, 24, 48 and 72 hours post-inoculation. The virus titers were determined by plaque assay in MDCK cells. Ten-day-old chicken embryonated eggs were infected with 100 plaque-forming units (100PFU/egg) of wild-type and mutant viruses of UDL-01/08 AIVs. viral replication efficiency in chicken embryonated eggs was determined up to 48 hrs post virus inoculation. The virus titres was determined using plaque assay in MDCK cells (C). One-way ANOVA with multiple comparisons was used to compare virus titres from each time point. In MDCK and CK cells, coloured asterisk next to a same-colour curve indicates that that virus has a significantly different titre from the remaining viruses and coloured asterisk next to a different-colour curve indicates a statistical difference in titre between those two viruses (* equals P < 0.05,** equals P < 0.01, *** equals P < 0.001, **** equals P < 0.0001).

The analysis of replication competency of these mutant viruses in CKCs also revealed that the mutant virus with the positively charged amino acid K at position 149 (G149K) in the HA showed replication patterns similar to the wt-UDL-01/08 virus at all post-infection time points in CKCs. The mutant virus with the negatively charged amino acid D or E at position 149 (G149D, G149E) in the HA replicated at higher rate with approximately 10-fold increase in viral titre compared to both the wt-UDL-01/08 virus and the mutant virus with the positively charged residue G149K at HA across all time points from 12 to 72 hrs post-infection (Figure 4B).

The impact of G149E mutation in the HA on H9N2 virus replication competency in chicken embryos was also analysed. Ten-day old hens’s embryos were inoculated with 100 PFU of the wt-UDL-01/08 virus or mutant viruses carrying G149E, G149D, or G149K mutations. After 48 hours, the viral titres were determined using the plaque assay. The results revealed that both the G149E and G149D mutant viruses replicated to levels comparable to the wt-UDL-01/08 virus (∼ 10^8^ PFU/ml), reflecting patterns observed in CKCs. In contrast, the G149K mutant replicated significantly lower than both the wt-UDL-01/08 virus and the mutants with the G149E and G149D substitutions (Figure 4C).

Altogether, the results conclude that the mutant viruses with negatively charged residues D or E at the HA position 149 exhibited replication fitness comparable to or greater than the wt-UDL-01/08 virus. Conversely, the mutant virus with the positively charged amino acid lysine (K) at the same position displayed reduced replication capabilities when compared to the mutant viruses with negatively charged amino acids at position 149 in the HA, both in cultured cells and in chicken embryos.

### The G149E mutant virus formed plaques larger in size compared to those of the wt-virus

The impact of the G149E mutation on virus plaque morphology was examined. Mutant viruses with negatively charged amino acid mutations (G149E and G149D) in the HA produced larger plaques compared to the wt-UDL-01/08 virus. In contrast, the mutant virus with the positively charged amino acid substitution (G149K) produced plaques with similar size with the wt-UDL-01/08 virus (Figure 5). The data suggests that the negatively charged amino acid at position 149 likely facilitates optimal virus-host cell surface interactions, allowing efficient cell-to-cell spread. On the other hand, a positive charge potentially leads to a strong attraction with cell surface receptors, restricting the formation of smaller plaques. Notably, the variations in plaque size and morphology did not correlate with the virus replication efficiency in MDCK cells. Altogether, this suggests that mutations at position 149 in the HA influence the biological properties of the H9N2 virus on multiple levels.

**Figure 5.**
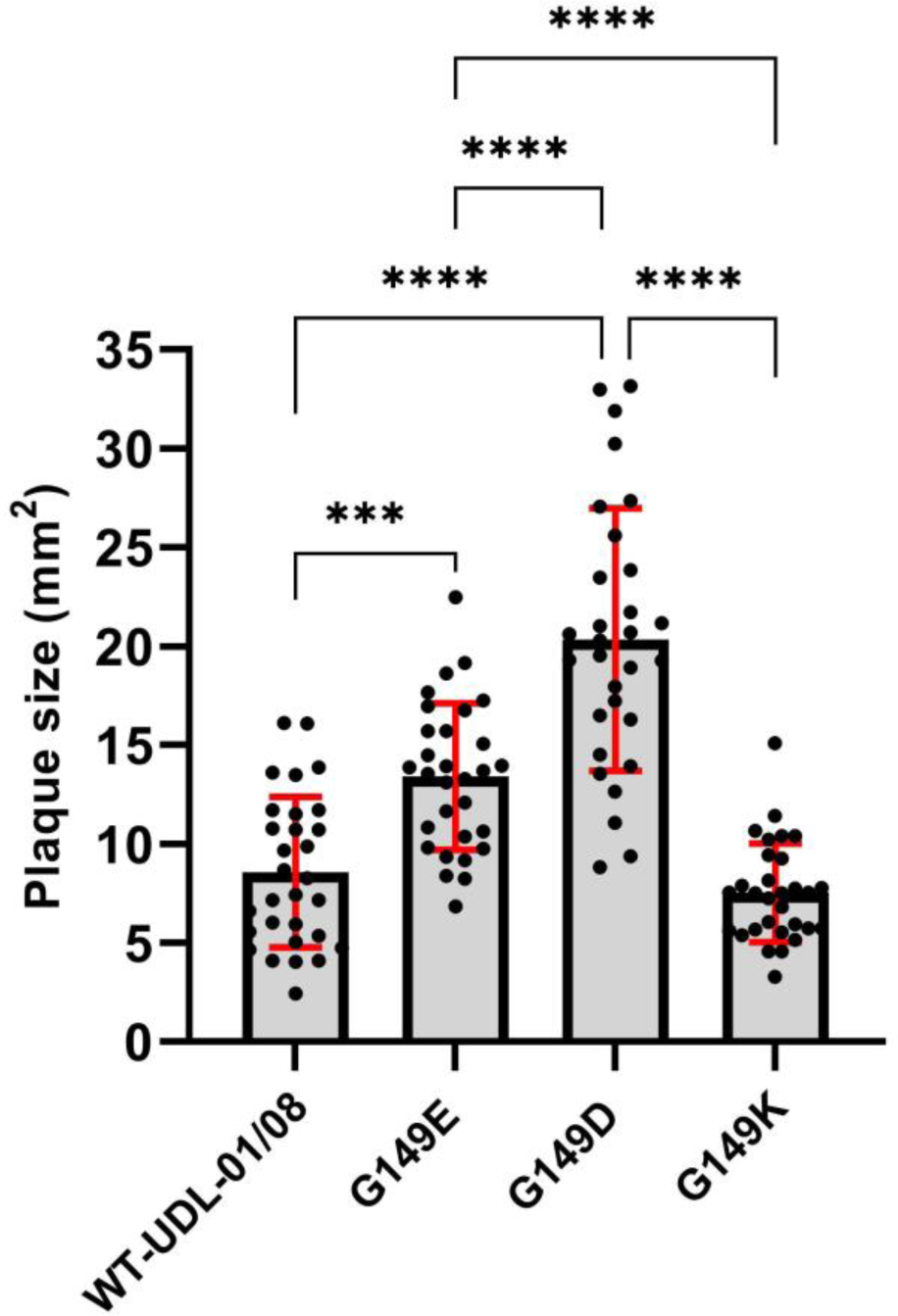
Plaque morphology of wt-UDL-01/08 and mutant virus carrying G149E, G149D and G149K mutation in MDCK cells. The plaque assay was performed in MDCK cells with wt-UDL-01/08 virus and mutant viruses with indicated amino acid substitutions at position 149 (G149E, G149D and G149K). D and E are negatively charged amino acid and K is a positively charged amino acid. Thirty plaques were randomly selected from each virus to measure the plaque size. ImageJ software was used to measure plaque size. One-way ANOVA was used to compare the plaque size of viruses between each viruses. Results shown are representative of three experimental results. Each black-coloured dot indicate a single plaque. Error bar = standard deviation. ***, P<0.001; ****, P<0.0001. G-Glycine, D-Aspartic acid, E-glutamic acid, K-lysine. mm2 – squire millimetres.

## Discussion

AIVs acquire mutations to maintain their fitness within the target host species. To pinpoint the amino acid residues susceptible to evolution due to immune pressure, we cultured the H9N2 (wt-UDL-01/08) virus in cells supplemented with antisera from chickens vaccinated with the homologous virus antigen. The progeny viruses recovered from immune escape experiments contained a G149E substitution. Haemagglutinin assays revealed that the mutant G149E-UDL-01/08 virus lost its ability to agglutinate chicken RBCs. Since the haemagglutination assays and haemagglutination inhibition (HI) assays are employed as diagnostic tools for H9N2 virus surveillance and antigenic characterisation (29, 41),the emergence of variants with G149E substitution potentially compromise the effectiveness of these assays for both surveillance and diagnostic purposes. The loss of haemagglutination activity due to mutations in viral HA of other influenza virus subtypes, such as human influenza A (H1N1) and (H3N2) viruses, has also been reported (31, 35). In H1N1 influenza virus, the substitution at position 225 (G225D) was responsible for the loss of haemagglutination activity, while in H3N2 virus, an amino acid change from Glu to Asp at position 190 (E190D) had the same effect (31, 34, 35).The viral surface glycoprotein HA is able to bind to the sialic acid receptor molecules present on host cells surface is governed by electrostatic interactions. Changes in amino acid polarity (neutral, negative or positive) due to mutations alter the overall electrostatic balance, disrupting viral binding affinity to the receptors on cell surface. The loss of agglutination in the mutant G149E-UDL-01/08 virus was due to the substitution of amino acid G to E at position 149. Amino acid G is a neutral (non-polar) residue, whereas amino acid E has negative polarity. Thus, the G149E substitution resulted in a change in polarity from neutral to negative, might be critical for the virus’s ability to bind to RBCs and cause agglutination. To test this hypothesis, we generated mutant viruses with changes at position 149, replacing amino acid G with either negatively charged amino acid D or positively charged amino acids K, R or H. The results provided clear evidence that, like the negatively charged amino acid D substitution, the impact was similarly significant to the G149E substitution of mutant G149E-UDL-01/08 virus. Negatively charged amino acid at position 149 possibly altered the electrostatic balance between the virus and RBC interaction, resulting in a loss of virus agglutination activity. Other studies have also reported similar observations, where the substitution of a negatively charged amino acid from G to D and E to D affected the RBCs agglutination activity of H1N1 and H3N2 influenza viruses, respectively (31, 34). On the other hand, all mutant viruses carrying positively charged amino acids (K/R/H) at position 149 retained haemagglutination activity.

To demonstrate the function of amino acid polarity at position 149 in contribution to receptor binding affinity, we modelled different sequences of wt-UDL-01/08 HA with amino acids G, E or K at position 149 using SWISS-MODEL (http://swissmodel.expasy.org). The charge distribution of all single amino acid substitutions at position 149 (G, E or K) were mapped onto H9HA structure using PyMOL software. The charge distribution difference in RBS of H9HA highlighted that change of amino acid from G to E increased negative charge distribution in and around the RBS, specially towards the 190-helix. While amino acid substitution from G to K at the same position increased positive charge distribution within and near RBS. The change in charge distribution in RBS has the most apparent effect on the binding of the ligand to the pocket of RBS. Positively charged RBS anchors to sialic acid molecules with greater strength, while negatively charged RBS does not bind with the same strength. In agreement with charge distribution differences, receptor binding results showed that the mutant G149E-UDL-01/08 virus carrying G149E mutation in HA decreased (> 10-fold) in 3SLN(6su) (sulfated variant of the classical avian-like receptor analogue) binding, while lost binding to 6SLN (human-like receptor analogue) binding compared to wt-UDL-01/08 virus. This demonstrated that charge of RBS correlates well with receptor binding and therefore, this may be the one of the reasons why HA of the mutant G149E-UDL-01/08 virus carrying G149E mutation does not agglutinate chicken erythrocytes. Further dissection of RBS of H9HA of H9N2 viruses indicated that 220-loop of RBS is involved in agglutination of chicken erythrocytes and residues at 215, 216 and 217 in 220-loop of RBS of H9HA of wt-UDL-01/08 virus is equally important for agglutination of chicken erythrocytes. More importantly, the mutant G149E-UDL-01/08 virus produced comparable progeny viruses to the wt-UDL-01/08 virus in mammalian and avian cells with larger plaque size compared to wt-UDL-01/08 virus.

The key point regarding the G149E mutation in the H9HA protein is that the mutant G149E-UDL-01/08 virus maintained stability without acquiring any additional compensatory substitutions in the HA or NA glycoproteins, even after being propagated multiple times in chicken embryos. Additionally, the introduction of the G149E mutation into two distantly related H9N2 viruses confirmed that the G149E substitution can maintained its fitness across different strains of H9N2 viruses. The observed data suggest that H9N2 viruses can potentially acquire the G149E mutation in the field, giving them the capability to cause sustained infections in poultry flocks. Given the proven genetic reassortment capability of H9N2 with other AIVs, the persistence of such viruses which are not detectable in the environment using routine assays might increase the likelihood of them donating their internal gene segments to other viruses. This could, in turn, enhance the zoonotic potential of the recipient AIV subtypes (4, 19-22).

The mutant G149E-UDL-01/08 virus had reduced antigenic cross-reactivity with the polyclonal antisera raised against the wt-UDL 01/08 virus. This suggested that the 149 position is a critical antigenic site in the HA of H9N2 viruses. Published data further indicate that mutations at the H9HA residue 149 decrease the HI titers of anti-H9HA monoclonal antibodies, confirming that this residue likely defines the antigenic site of H9N2 viruses (42). Since the prevention and control of H9N2 infections in poultry in endemic regions are based on vaccination (43, 44), viruses with G149E mutations can potentially evade the immunity administrated with current vaccines. Moreover, the mutant G149E-UDL-01/08 virus cannot be detected by routine haemagglutination assays employed in surveillance programs, particularly in resource-limited laboratories, since qPCR detection is more costly. Therefore, the emergence of the mutant G149E-UDL-01/08 virus may poses a continued threat to poultry and a risk of zoonotic infections.

The results conclude that both the haemagglutination assays and HI assays are essential for avian influenza surveillance and for characterising the antigenic properties of the HA protein, especially when selecting vaccine seeds in resource-limited laboratories. The emergence of variants in the field with G149E mutations suggests that they might not be neutralized by the immunity conferred by current vaccines and may also evade detection by ongoing surveillance programs. Consequently, these mutants could remain undetected in the field, potentially giving rise to novel viruses by donating their internal gene segments, which can lead to new phenotypic traits with zoonotic or pandemic potential. This highlights the urgent need for continuous monitoring of H9N2 and co-circulating subtypes in avian populations.

## Materials and Methods

### Ethics Statement

All the procedures involving embryonated eggs were carried out in strict accordance with the guidance and regulations of United Kingdom Home Office regulations under project licence number P68D44CF4. As part of this process, the work has undergone scrutiny and approval by the animal welfare ethical review board at the Pirbright Institute. All the influenza virus related work was carried out under enhanced biosafety containment level-2 laboratory and animal experimental facilities.

### Cells

Chicken Kidney (CK) cells were prepared three days prior to use in virus growth kinetic assays in 12-well plates as previously described (45) and maintained in CK cell media comprised of Eagle’s Minimum Essential Medium (EMEM) (Sigma-Aldrich) supplemented with 2 mM L-glutamine, 0.6% w/v Bovine Serum Albumin (BSA), 1% Penicillin–Streptomycin (PS) and 10% v/v Tryptose Phosphate Broth (TPB). Madin Darby canine kidney (MDCK) cells and human embryonic kidney 293T cells (ATCC) were maintained in Dulbecco’s Modified Eagle’s Medium (DMEM) (Life Technologies Thermo Fisher) supplemented with 10% (v/v) Foetal Calf Serum (FCS, Gibco) and Penicillin (100U/ml), Streptomycin (100µg/ml) (Thermo Fisher Scientific) at 37 °C with 5% CO_2_.

### Influenza A viruses prepared by reverse genetics and their propagation in eggs

The nucleotide sequences of all gene segments of H9N2 viruses were access and retrieved from publicly accessible databases, namely the Global Initiative on Sharing All Influenza Data (GISAID https://www.gisaid.org/) or the National Center for Biotechnology Information (NCBI https://www.ncbi.nlm.nih.gov/). Gene segments were synthesised by Geneart^TM^ (Thermo-Fisher Scientific) and subcloned into the pHW2000 vector. The three H9N2 influenza viruses used in the study namely, A/chicken/Pakistan/UDL-01/2008 (NCBI Accession number.CY038455), A/Environment/Bangladesh/26218/2015 (NCBI accession no. KY635657), and A/chicken/Vietnam/H7F-14-BN4-315/2015 (GISAID Accession number. EPI_ISL_327772). All three wild-type viruses and their mutants’ viruses were rescued by reverse genetic (46) and subsequent propagation in 10-day-old specific pathogen free (SPF) embryonated chicken eggs at 37 °C for 72 hours.

### Site-directed mutagenesis

The QuickChange Lightning Kit (Agilent) was used for site-directed mutagenesis in HA gene for residues 149, 183, 186, 188, 215, 216, 217. The HA and NA gene sequences of wild-type and mutant viruses were confirmed by sanger sequencing at Source Bioscience (Cambridge, UK).

### Haemagglutination assay

The HA assays were performed with 1% chicken RBCs. Briefly, 50µl of virus sample was serially diluted in PBS in two-fold in V-bottom 96-well plate (Greiner) and followed by adding 50µl of 1% washed chicken RBCs. The plates were incubated for 30 minutes at room temperature before the HA titre was recorded and presented as HA units (HAU) per 50µl.

### Virus plaque Assay

Virus samples were serially diluted in serum free Dulbecco’s modified Eagle’s medium (SF-DMEM) and 200μl from each dilution was overlayed in duplicate on MDCK cells in 12-well tissue-culture plates. After 1 hour incubation at 37 °C, 5% CO2, inoculum was removed, cells were washed with PBS and then cells were overlayed with plaque assay overlay medium containing 2 μg/ml TPCK trypsin and 0.6% agar. After 72 hours incubation at 37 °C, 5% CO2, agar plugs were removed, and cells were fixed and stained with crystal violet. The number of plaque forming units (PFU) were counted and accordingly PFU/ml were calculated. The plaque images were taken by Panasonic camera FZ38 and randomly selected 30 plaques from each virus were analysed for plaque size.

### Virus microneutralization assay

Virus neutralization assays were performed as previously described (47). Briefly, heat-inactivated antisera were serially diluted and 100 TCID_50_ of virus was added to each dilution of the dilution series of heat-inactivated antisera and incubated for 1 hour at 37 °C. Virus/antisera mix was added to confluent MDCK cells and incubated for 1 hour at 37 °C. Thereafter, cells were washed with PBS and overlayed with SF-DMEM containing 2μg/ml TPCK trypsin. 72 hours post-infection, medium was removed, cells were fixed and stained with crystal violet before serum neutralization titres were determined.

### Virus growth curves

Growth kinetics of the wild-type and mutant viruses were assessed in MDCK cells and primary chicken kidney (CK) cells. Briefly, MDCK and primary chicken kidney (CK) cells were infected in triplicate with wild-type and mutant viruses at a MOI 0.001 and 0.01, respectively for 1h at 37 °C. Then cells were washed twice with phosphate-buffered saline (PBS) to remove unbound virus and 2 ml of virus growth medium (DMEM plus 2 μg/ml tosyl phenylalanyl chloromethyl ketone [TPCK]-treated trypsin for MDCK cells and Eagle’s minimum essential medium [EMEM], 0.6% bovine serum albumin [BSA], and 10% tryptose phosphate broth [TPB] for CKCs) was added. The cell supernatant was harvested after 12-, 24-, 48- and 72-hours post infection, and virus titers were estimated in triplicate by using plaque assay in MDCK cell.

### *In-ovo* growth

Ten-day-old chicken embryonated eggs were infected with 100 plaque-forming units of wild-type and mutant viruses of UDL-01/08 AIVs. The eggs were incubated at 37 °C for 48 hrs before being chilled at 4°C. Allantoic fluid was harvested and titrated by haemagglutination (HA) assay and plaque assay.

### Passage of virus in eggs

Virus samples were serially diluted in 10-fold in PBS and each dilution was added to a single egg. Eggs were harvested 72 hrs post-infection and tested for presence of virus by haemagglutination assay. For each virus, the highest dilution with positive HA titre was continuously further passaged for 5 times and sequenced.

### Sequencing of viruses

Sequencing of virus HA and NA genes was performed using sanger sequencing. Briefly, RNA from viruses was extracted from egg allantoic fluid and cell culture medium by using the QIAamp viral RNA minikit (Qiagen). A Verso cDNA synthesis kit (Thermo Scientific) was used to perform reverse transcription with the universal FluA primer. PCR was undertaken by using primers specific for HA1 and HA2 adjunct with 5’ M13F and M13R sequencing motifs. The QIAquick PCR purification kit (Qiagen) was used to purify the resulting products, which were then sequenced by using the M13F and M13R sequencing primers.

### Virus purification

Embryonated chicken eggs were infected with virus and virus from clarified egg allantoic fluid was initially pelleted by ultracentrifugation at 27,000 rpm for 2 h. Then pellets were resuspended in PBS and before homogenized using a glass homogenizer. Homogenized virus samples were purified by using a continuous 30% to 60% (wt/vol) sucrose gradient (48). The concentration of purified virus samples was determined by enzyme-linked immunosorbent assay against NP as prescribed previously (33).

### Biolayer interferometry

Binding of virus and sialic receptor analogous was measured using an Octet Red biolayer interferometer (Pall FortéBio) as previously described (37). Briefly, receptor analogous of sialoglycopolymers containing polyacrylamide backbone conjugated to three different types of trisaccharide α2,6-sialyllactosamine (6SLN), α2,3-sialyllactosamine (3SLN), or Neu5Ac α-2,3Gal β1-4(6-HSO_3_) GlcNAc (3SLN(6su)) and biotin (Lectinity Holdings) were used. Sialylglycopolymers were immobilized on streptavidin-coated biosensors (Pall FortéBio) at concentrations ranging from 0.01 to 0.5 µg/ml in a solution containing 10 mM HEPES (pH 7.4), 150 mM NaCl, 3 mM EDTA, and 0.005% Tween 20 (HBS-EP). Virus was diluted in HBS-EP buffer (Teknova) containing 10 μM oseltamivir carboxylate (Roche) and 10 µM zanamivir (GSK) to a concentration of 100 pM. The association between virus and immobilized receptors was measured at 20 °C for 30 minutes. Virus binding amplitudes were normalized to fractional saturation and plotted as a function of sugar loading. Virus relative estimated dissociation constants (KDs) were calculated as described in previous (37).

### Modelling of haemagglutinin structures and charge distribution

Structure prediction and modelling of H9HA and derivative mutations were performed using SWISS-MODEL (http://swissmodel.expasy.org) homology modelling server. Images of structure of all wild-type and mutant H9HA proteins and their charge distribution were generated by PyMol version 4.6 (www.pymol.org).

### Statistical analysis

Statistical analyses were performed using GraphPad Prism 8 (GraphPad Software). One-way ANOVA and Tukey’s multiple comparison test was used to analyse the differences between different groups. p values < 0.05 were considered significant.

## Acknowledgements

The authors extend their deep gratitude to Dr. Daniel H. Goldhill from the Royal Veterinary College, London, for providing invaluable guidance on this work. Additionally, we would like to express our sincere appreciation to Dr. Thomas Peacock for conducting a thorough review of this manuscript. *

---

**Supplementary Figure 1.**
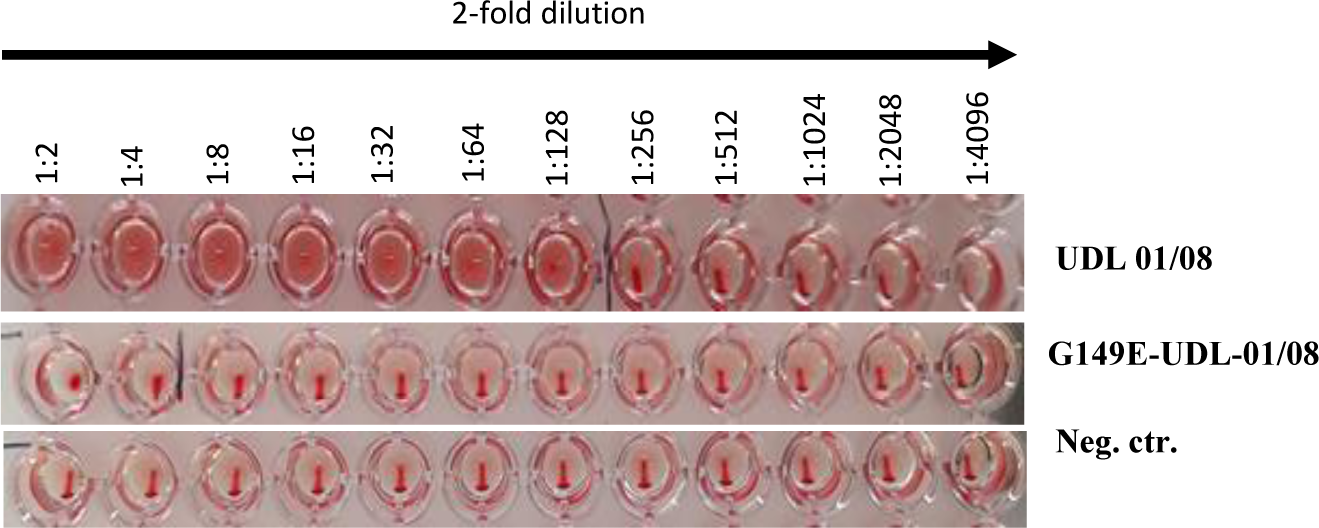
Analysis of haemagglutination of chicken RBCs with wt-UDL 01/08 and G149E-UDL-01/08 mutant viruses. 96-well plate containing 1% chicken RBCs in line 1 were treated with wt-UDL-01/08, line 2 were treated with G149E-UDL 01/08 mutant virus and Line 3 were mock treated with PBS as negative control (Neg. ctr.) for haemagglutination test. Both wt-UDL-01/08 and G149E-UDL-01/08 mutant viruses were propagated in embryonated chicken eggs.

**Supplementary Figure 2.**
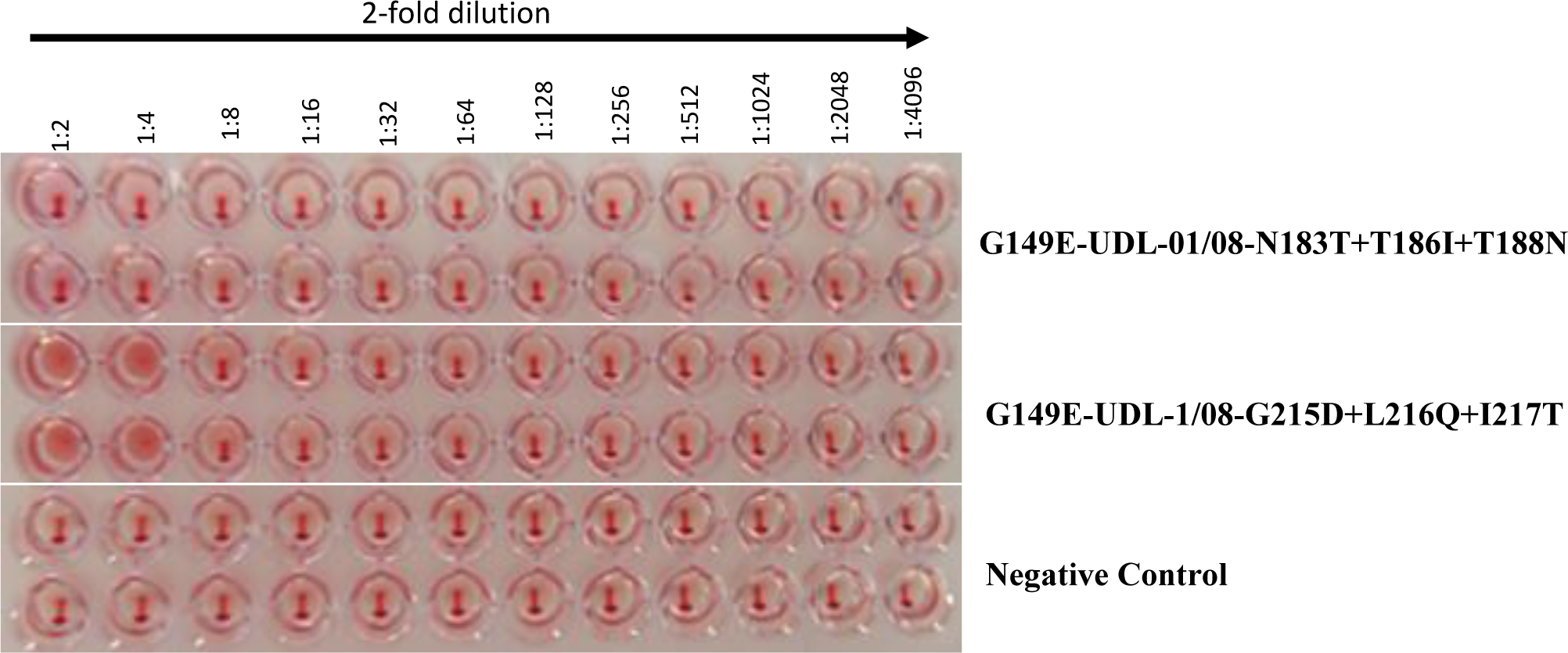
Analysis of haemagglutination of chicken RBCs with two UDL-01/08 mutant viruses converted their RBS (190-helix and 220-loop of RBS) separately into BD/26218/15 virus RBS while carrying HA G149E mutation. 96-well plate containing 1% chicken RBCs in line 1 were treated with G149E-UDL-01/08-N183T+T186I+T188N mutant virus, line 2 were treated with G149E-UDL 1/08-G215D+L216Q+I217T mutant virus and Line 3 were mock treated with PBS as negative control (Neg. ctr.) for haemagglutination test. Both mutant viruses were propagated in embryonated chicken eggs. Haemagglutination assays of two mutant viruses were performed with the virus samples with same PFU/ml for standardize number of live virus particles in virus samples. N-asparagine, T-threonine, I-isoleucine, G-glycine, D-aspartic acid, L-leucine, and Q-glutamine.

**Supplementary Figure 3.**
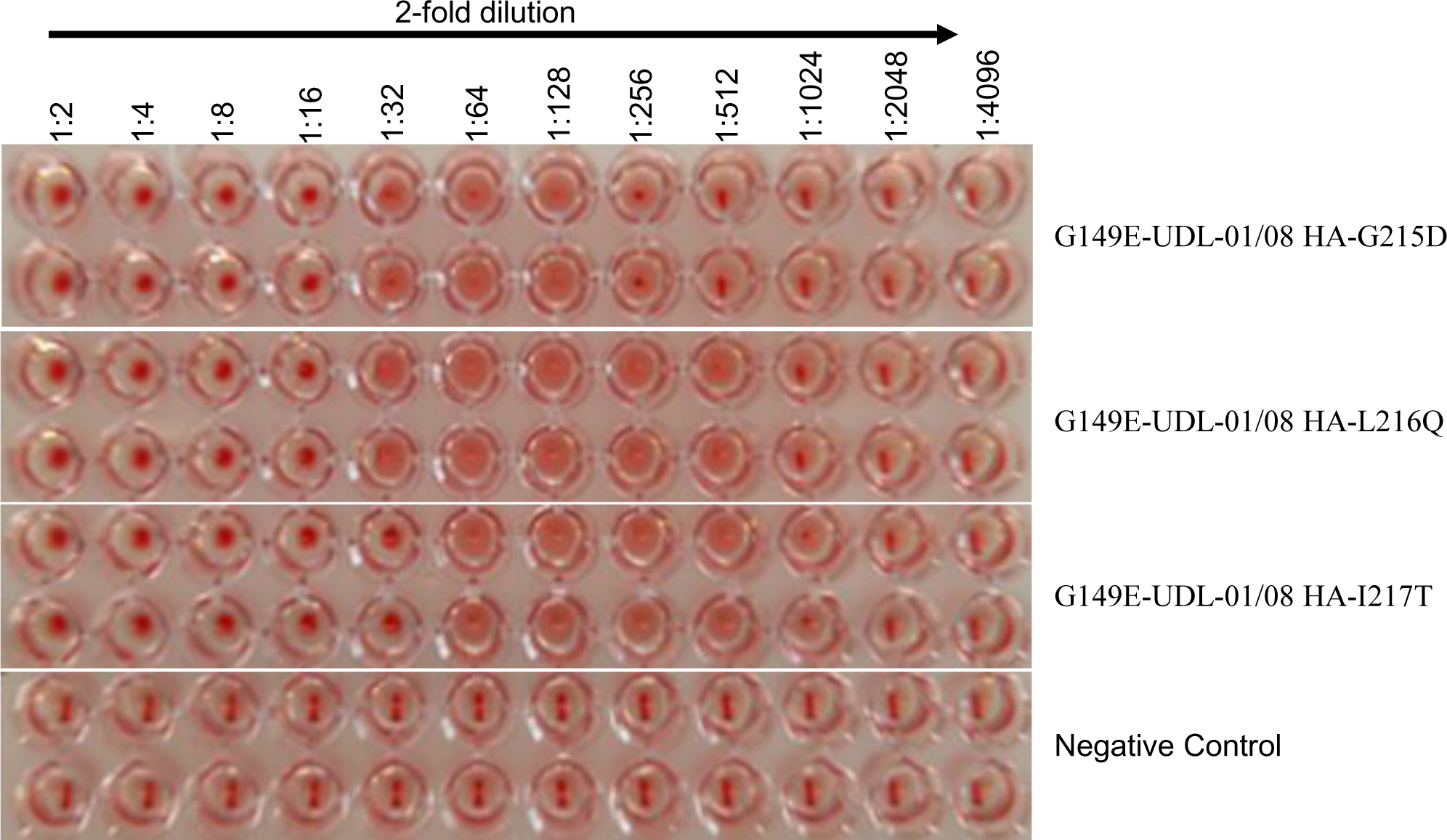
Analysis of haemagglutination of chicken RBCs with UDL-01/08 mutant viruses carrying G149E substitution with converted position at 215, 216 and 217 in 220-loop of RBS separately into BD/26218/15 virus RBS. 96-well plate containing 1% chicken RBCs in line 1 were treated with G149E-UDL-01/08-G215D mutant virus, line 2 were treated with G149E-UDL 1/08-L216Q mutant virus, Line 3 treated with G149E-UDL 1/08-I217L mutant virus and line 4 were mock treated with PBS as negative control (Neg. ctr.) for haemagglutination test. All mutant viruses were propagated in embryonated chicken eggs. T-threonine, I-isoleucine, G-glycine, D-aspartic acid, L-leucine, and Q-glutamine.

**Supplementary Figure 4.**
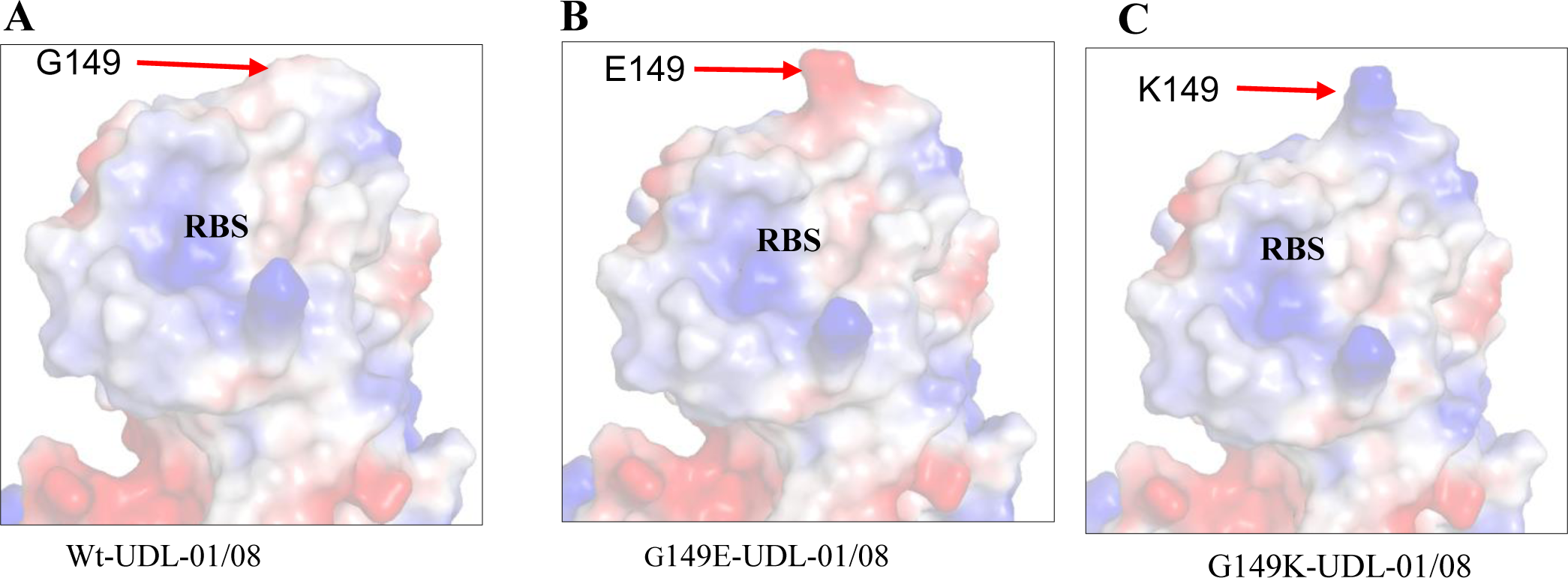
Distribution of surface charges around the receptor binding site (RBS) on the UDL-01/08 virus haemagglutinin (HA) protein. A) wt-UDL-01/08, B) G149E-UDL-01/08, C) G149K-UDL-01/08. Blue and red colors represent basic and acidic charges, respectively. RBS-receptor binding site. G-glycine, E-glutamic acid and K-lysine. **

