## Supplemental figures for "Investigation of H9N2 avian influenza immune escape mutant that lacks haemagglutination activity"

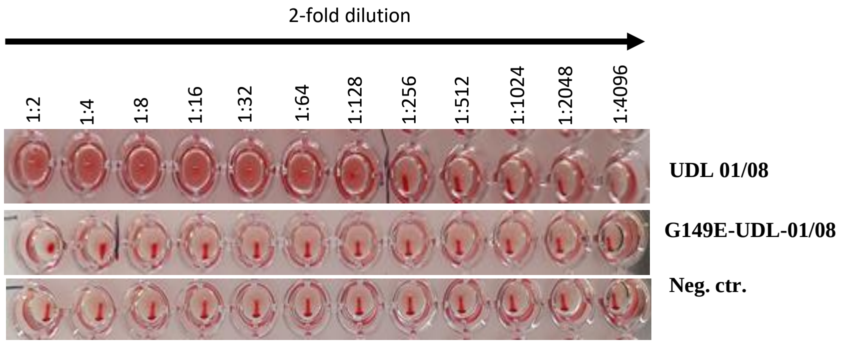


**Supplementary Figure 1. Analysis of haemagglutination of chicken RBCs with wt-UDL 01/08 and G149E-UDL-01/08 mutant viruses.** 96-well plate containing 1% chicken RBCs in line 1 were treated with wt-UDL-01/08, line 2 were treated with G149E-UDL 01/08 mutant virus and Line 3 were mock treated with PBS as negative control (Neg. ctr.) for haemagglutination test. Both wt-UDL-01/08 and G149E-UDL-01/08 mutant viruses were propagated in embryonated chicken eggs.

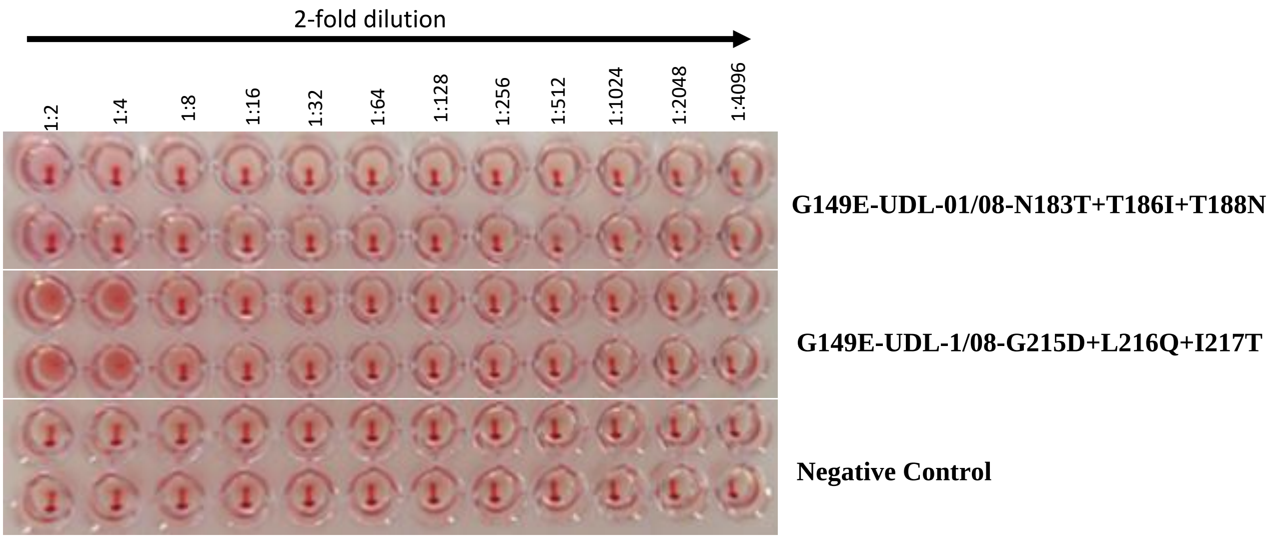


**Supplementary Figure 2. Analysis of haemagglutination of chicken RBCs with two UDL-01/08 mutant viruses converted their RBS (190-helix and 220-loop of RBS) separately into BD/26218/15 virus RBS while carrying HA G149E mutation**. 96-well plate containing 1% chicken RBCs in line 1 were treated with G149E-UDL-01/08-N183T+T186I+T188N mutant virus, line 2 were treated with G149E-UDL 1/08-G215D+L216Q+I217T mutant virus and Line 3 were mock treated with PBS as negative control (Neg. ctr.) for haemagglutination test. Both mutant viruses were propagated in embryonated chicken eggs. Haemagglutination assays of two mutant viruses were performed with the virus samples with same PFU/ml for standardize number of live virus particles in virus samples. N-asparagine, T- threonine, I-isoleucine, G-glycine, D-aspartic acid, L-leucine, and Q-glutamine.

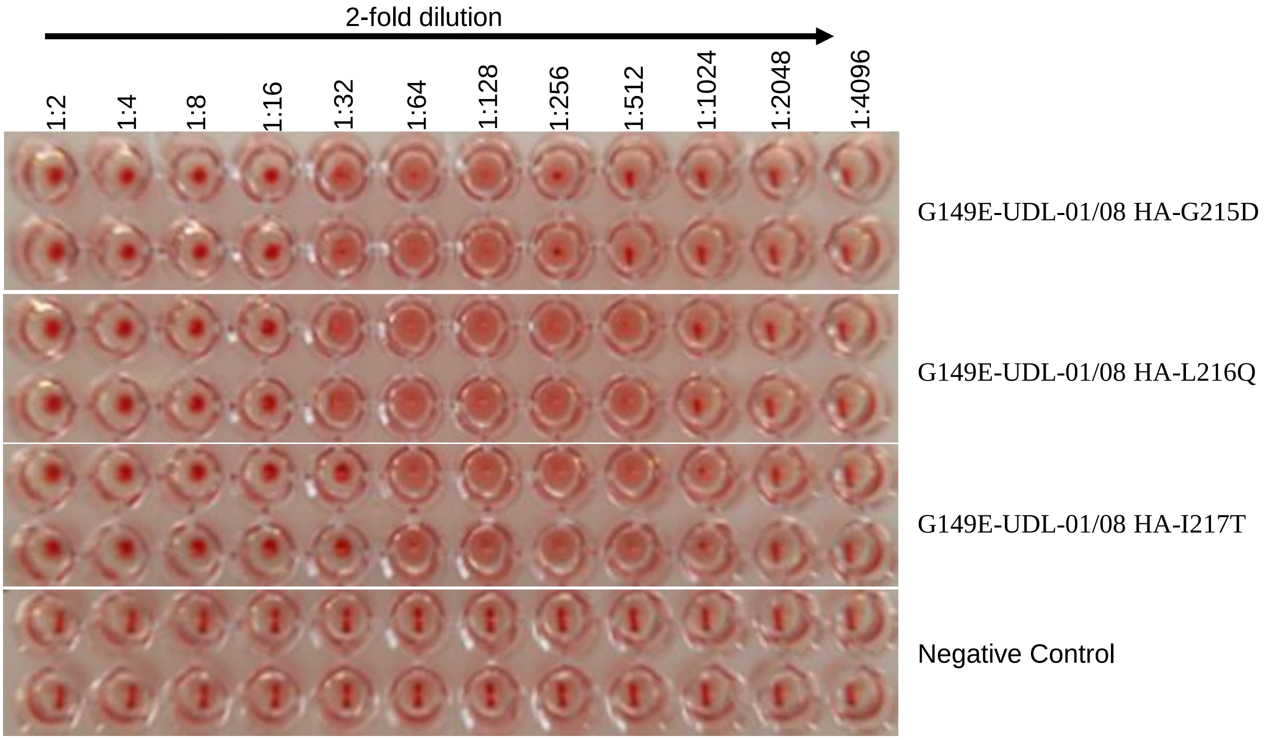


**Supplementary Figure 3. Analysis of haemagglutination of chicken RBCs with UDL-01/08 mutant viruses carrying G149E substitution with converted position at 215, 216 and 217 in 220-loop of RBS separately into BD/26218/15 virus RBS.** 96-well plate containing 1% chicken RBCs in line 1 were treated with G149E-UDL-01/08-G215D mutant virus, line 2 were treated with G149E-UDL 1/08-L216Q mutant virus, Line 3 treated with G149E-UDL 1/08-I217L mutant virus and line 4 were mock treated with PBS as negative control (Neg. ctr.) for haemagglutination test. All mutant viruses were propagated in embryonated chicken eggs. T- threonine, I-isoleucine, G-glycine, D-aspartic acid, L-leucine, and Q-glutamine.

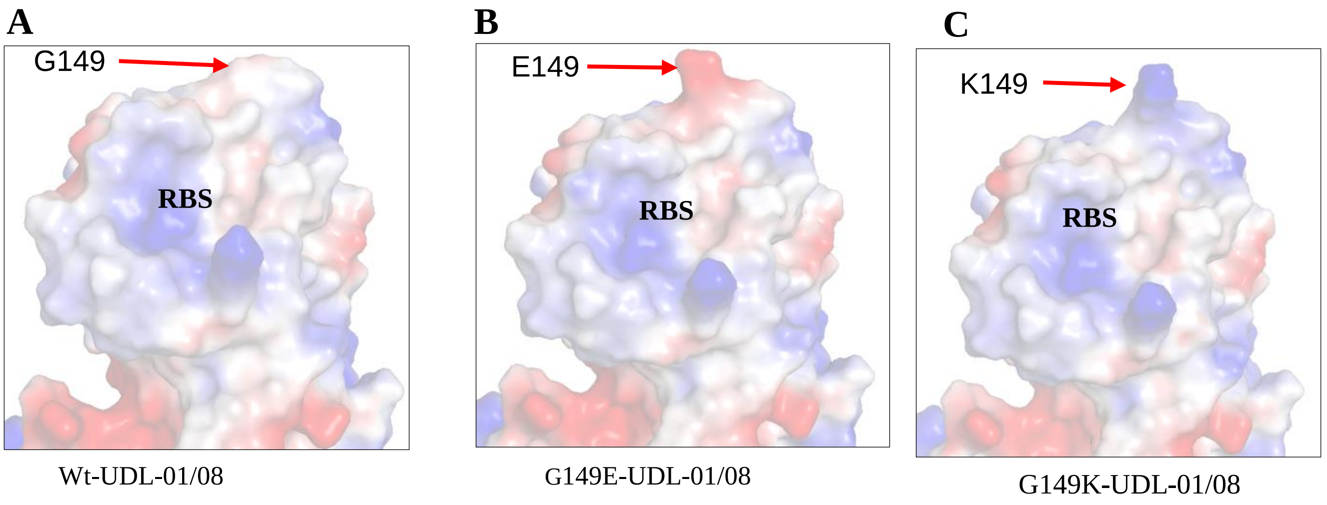


**Supplementary Figure 4. Distribution of surface charges around the receptor binding site (RBS) on the UDL-01/08 virus haemagglutinin (HA) protein.** A) wt-UDL-01/08, B) G149E-UDL-01/08, C) G149K-UDL-01/08**.** Blue and red colors represent basic and acidic charges, respectively. RBS-receptor binding site. G-glycine, E-glutamic acid and K-lysine. **
